# VGF in cerebrospinal fluid, when combined with conventional biomarkers, enhances prediction of conversion from mild cognitive impairment to Alzheimer’s Disease

**DOI:** 10.1101/512939

**Authors:** Daniel A. Llano, Priya Devanarayan, Viswanath Devanarayan, for the Alzheimer’s Disease Neuroimaging Initiative (ADNI)

## Abstract

Sensitive and accurate biomarkers for the prediction of conversion from mild cognitive impairment (MCI) to Alzheimer’s Disease (AD) are needed to both support clinical care and enhance clinical trial design. Here, we examined the potential of cerebrospinal fluid (CSF) levels of a peptide derived from a neural protein involved in synaptic transmission, VGF (a non-initialism), to enhance accuracy of prediction of conversion from MCI to AD. The performance of conventional biomarkers (CSF Aβ1-42 and phosphorylated tau +/− hippocampal volume) was compared to the same biomarkers with CSF VGF peptide levels. It was observed that VGF peptides are lowered in patients with AD compared to controls and that combinations of CSF Aβ1-42 and phosphorylated tau, hippocampal volume and VGF peptide levels outperformed conventional biomarkers alone (hazard ratio = 2.2 vs. 3.9). VGF peptide levels were correlated most strongly with total tau levels, but not hippocampal volume, suggesting that they serve as a marker for neuronal degradation, but not necessarily in the hippocampus. The latter point suggests that VGF may serve as a more general marker of neurodegeneration. Future work will be needed to determine the specificity of VGF for AD vs. other neurodegenerative diseases.

## Introduction

Alzheimer Disease (AD) is characterized by a long prodromal course during which a number of pathological changes occur prior to the onset of clinical symptoms. Classically, these changes include the deposition of amyloid beta (Aβ) and phosphorylated tau (pTau) into the brain, hippocampal atrophy and disruptions of metabolism, particularly in the temporal and parietal cortices (for review of preclinical pathology and biomarkers, please see [1]). It is speculated that these biomarkers are part of a cascade whereby Aβ triggers a series of pathological events, leading to neuronal dysfunction, hyperphosphorylation of tau and consequent synaptic loss, leading to volume loss and metabolic disruption [2–4]. These changes have formed the basis for the use of a series of fluid and imaging biomarkers to facilitate clinical and research practice.

AD biomarkers may be used to 1) achieve earlier diagnoses for patients, 2) predict which individuals are most likely to clinically worsen over time, 3) help to identify and stratify subjects enrolling in AD-related clinical trials and 4) serve as outcome measurements in AD-related clinical trials [5–7]. For example, there is a 10-15% misdiagnosis rate when AD is diagnosed on clinical grounds only. This high rate of misdiagnosis has substantial cost implications [8–11] and if such misdiagnosed subjects are enrolled into clinical trials, they could obscure the impact of disease-modifying therapy. In addition, prediction of clinical decline in subjects with early-stage disease will permit the institution of aggressive interventions, such as physical exercise or pharmacologic therapy, to stave off AD symptoms. Finally, novel biomarkers or combinations of biomarkers could be used to enrich MCI clinical trials with subjects with high conversion rates to shorten and diminish the cost of clinical trials [12, 13]. Therefore, a better understanding of how biomarkers delineate disease classes and predict progression is needed.

Recently, our group and others have identified a group of novel plasma and cerebrospinal fluid (CSF) biomarkers that fall outside the traditional Aβ cascade. Many of these markers have been shown to be useful in the prediction of MCI to AD conversion [14–20]. For example, we used a hypothesis-free bioinformatics approach to identify a panel of 16 peptides in CSF initially identified as showing high diagnostic accuracy for AD vs. control, that was highly predictive of conversion from mild cognitive impairment (MCI) to AD in an independent group of subjects and outperformed conventional CSF markers such as Aβ, tau derivatives and their ratios [20]. These studies highlight non-canonical pathological cascades that may both provide useful tools for clinical practice and clinical trials purposes, and may also reveal new insights about disease mechanisms underlying AD.

One of the peptides identified using this hypothesis-free approach to separate AD from normal (NL) controls was VGF [20]. VGF (a non-initialism) has recently received significant attention because of its role in learning and memory and potential role in the pathophysiology of AD [21, 22]. VGF is a neurotrophin-inducible 615-amino acid polypeptide secreted by neurons and is cleaved into multiple smaller fragments ranging in length from 16-129 amino acids. VGF is produced in a number of brain regions, including the cerebral cortex, amygdala, hippocampus and hypothalamus, as well as in neuroendocrine tissues such as the adrenal medulla and adenohypophysis, and is thought to be involved in synaptogenesis and energy homeostasis [23, 24]. We and others have observed altered levels of VGF in the CSF of AD patients compared to controls, though not all studies had the same directionality (studies showing a decrease: [25–31], study showing an increase: [32]). VGF overexpression also protects against memory impairment in 5xFAD transgenic mice that model AD [21]. However, previous work has not yet examined the potential for VGF in the CSF, when combined with established biomarkers, to predict MCI to AD conversion. It should not be assumed that VGF independently contributes to the prediction of MCI to AD conversion: it is possible that it is a redundant marker for a process already encoded by changes in a more conventional biomarker.

Therefore, in the current study, we examined the potential for VGF in the CSF, when combined with conventional biomarkers of CSF Aβ1-42, total tau (tTau) and pTau-181 and hippocampal volume, to enhance the diagnostic and prognostic accuracy of these markers. The focus of this work is on the VGF peptide fragment with sequence NSEPQDEGELFQGVDPR (“VGF.NSEP”) since it previously emerged as a strong predictor in a panel of peptides that predict MCI to AD conversion [20], though other VGF peptide fragments are also examined. Unlike our previous studies involving hypothesis-free approaches to identify optimal peptides to include in biomarker signatures [20, 33, 34], the current study was focused on the utility of VGF. Using data from two independent groups in the ADNI cohort: one group of AD and control subjects and a separate group of MCI subjects, it was found that VGF, when combined with conventional biomarkers, enhanced both the diagnostic accuracy of these markers and the ability of these markers to predict MCI to AD conversion.

## Methods

Methods and data used for this research are similar to those used in Devanarayan et al. [33]. The ADNI database (adni.loni.usc.edu) utilized in this research was launched in 2003 as a public-private partnership, led by Principal Investigator Michael W. Weiner, MD. The primary goal of ADNI has been to test whether serial MRI, PET, other biological markers, and clinical and neuropsychological assessments can be combined to measure the progression of MCI and early AD. For up-to-date information, see www.adni-info.org. This study was conducted across multiple clinical sites and was approved by the Institutional Review Boards of all of the participating institutions. Informed written consent was obtained from all participants at each site. Data used for the analyses presented here were accessed on February 24, 2018. Although the ADNI database continues to be updated on an ongoing basis, most newly added biomarker data are from later time points (i.e., beyond 1 year), in contrast to the baseline data used in this study.

### Subjects

This research was focused on the relationship between VGF, conventional biomarkers (CSF amyloid/tau and MRI hippocampal volume [HV]) and therefore, only those subjects whose values for these markers were available at baseline were included in this study. Ultimately, this dataset included 287 subjects across the three diagnostic categories (AD, MCI and NL). NL subjects were defined as those without memory complaints and had a clinical dementia rating (CDR) score of 0. MCI subjects had CDR scores of 0.5, had an abnormal score on Wechsler Memory Scale Revised– Logical Memory II and did not have significant functional impairment. AD subjects had functional decline and a CDR score of 0.5 or 1.0.

### Hippocampal volume

HV was chosen given its robust ability to predict MCI to AD conversion [35, 36] and its incorporation into proposed schema to classify AD subjects [37]. HV was obtained from MRI scans (mostly 1.5T; 25% in this dataset had 3.0T scans) and was computed using FreeSurfer software. Please see “UCSF FreeSurfer Methods” PDF document under “MR Image Analysis” in the ADNI section of https://ida.loni.usc.edu/) for details as well as [38–40].

### CSF Samples

Innogenetics’ INNO-BIA AlzBio3 immunoassay on a Luminex xMAP platform (see [41] for details) was used to measure levels of the conventional biomarkers Aβ1-42, tTau, and pTau-181 in CSF. The Caprion Proteomics mass spectrometry platform was used to measure levels of individual peptides. The VGF peptides (sequence NSEPQDEGELFQGVDPR, referred to here as VGF.NSEP, sequence AYQGVAAPFPK, referred to here as VGF.AYQG and sequence THLGEALEPLSK, referred to here as VGF.THLG) used in this study were among a total of 320 peptides generated from tryptic digests of 143 proteins. Details regarding the measurements of these peptides can be found in the Use of Targeted Mass Spectrometry Proteomic Strategies to Identify CSF-Based Biomarkers in Alzheimer’s Disease Data Primer (found under Biomarkers Consortium CSF Proteomics MRM Data Primer at ida.loni.usc.edu) and in [19].

### Statistical Methods

As we have described previously [33], optimal combinatorial signatures including CSF Aβ1-42, tTau, pTau-181, their ratios, HV and VGF-derived peptides with simple decision thresholds for each marker were first identified from the AD and NL subjects. These signatures were revealed by an unbiased, data-driven manner via regression and tree-based computational algorithms called Patient Rule Induction Method [42] and Sequential BATTing [43]. To measure the performance of each signature for disease-state differentiation (i.e., NL vs. AD), five-fold cross-validation was performed. To do this, the data were randomly divided into five subgroups, referred to as folds, and a signature was derived from the remaining four folds. This signature was then tested on the left-out fold. This process was repeated for 10 iterations and median performance of each performance of positive predictive value (PPV), negative predictive value (NPV) and accuracy was computed.

Once an optimal signature for differentiating NL from AD was derived, it was tested on a different group of 135 MCI subjects from the ADNI dataset. Baseline values for Aβ1-42, tTau, pTau-181, HV and VGF peptides for each MCI subject at baseline were used to classify each subject as being “signature positive” (i.e., similar to the profile found in AD) or “signature negative” (i.e., similar to the profile found in NC). PPV, NPV and accuracy were then computed by comparing the actual outcome (conversion or not to AD over 36 months) to the predicted outcome (signature positive/negative which would predict conversion/nonconversion, respectively). Exact McNemar’s test was used to compare PPV, NPV and accuracy.

In addition to measuring the performance of whether MCI subjects would convert over 36 months, time to conversion was also computed using available data up to 10 years after the initial evaluation. Potential markers for this analysis were grouped into categories:

1. Demographic markers (presence of APO-E4 allele, age, gender, education)
2. Demographic markers + HV
3. Demographic markers + amyloid/tau CSF markers (heretofore called “AT”: Aβ1-42, tTau, pTau-181, ratios of tTau to Aβ1-42 & pTau-181 to Aβ1-42)
4. Demographic markers + HV + AT
5. Demographic markers + HV + AT + VGF

All analyses related to predictive modeling and signature derivation were carried out using R (http://www.R-project.org), version 3.4.1, with the publicly available package, SubgrpID [43]. The time to progression analysis of the derived signatures and related assessments were carried out using JMP®, version 13.2.

## Results

### Demographics

Similar to Devanarayan et al. (2019), 66 AD, 135 MCI and 86 NL subjects were included in the analysis and their demographic information and rates of conversion from MCI to AD are shown in Table 1. There were no statistically significant differences in terms of age or education (range of means = 75.1 to 75.8 years, p>0.05) and education (range of means = 15.1 to 16 years, p>0.05). There was a greater number of males than females (59.1 vs 40.9%), though their likelihood of conversion from MCI to AD over 36 months was similar (43.5% vs. 53.9%, p=0.285, Chi-squared test). The likelihood that an APO-E4 allele was present was higher AD than in other subjects (present in 71.2% AD, 50% MCI and 24.4% NL subjects, p < 0.0001, Chi-squared test) and was a relatively weak risk factor for the conversion of MCI to AD (present in 40/62 converters and 31/70 non-converters p=0.03, Chi-squared test), both of which have been demonstrated previously [44–46].

**Table 1:** Disease-state demographics

|  |  | AD | MCI | NL |
| --- | --- | --- | --- | --- |
|  |  | (n=66) | (n=135) | (n=86) |
| Gender (n) | M | 37 | 91 | 44 |
|  | F | 29 | 44 | 42 |
| Apo-E (n) | E4 | 47 | 71 | 21 |
|  | Non-E4 | 19 | 64 | 65 |
| Age<br>(years, Mean +/- SD) |  | 75.1 +/- 7.5 | 74.8 +/- 7.4 | 75.8 +/- 5.6 |
| Education<br>(years, Mean +/- SD) |  | 15.1 +/- 3 | 16 +/- 3 | 15.6 +/- 3 |
| MMSE<br>(Mean +/- SD) |  | 23.5 +/- 1.9 | 26.9 +/- 1.7 | 29.1 +/- 1 |

|  |  | MCI to AD<br>converters | Stable MCI |
| --- | --- | --- | --- |
|  |  | (n=64) | (n=71) |
| Gender (n) | M | 40 | 51 |
|  | F | 24 | 20 |
| Apo-E (n) | E4 | 40 | 31 |
|  | Non-E4 | 24 | 40 |
| Age<br>(years, Mean +/- SD) |  | 74.9 +/- 7.6 | 74.7 +/- 7.2 |
| Education<br>(years, Mean +/- SD) |  | 15.6 +/- 3.0 | 16.4 +/- 2.9 |
| MMSE<br>(Mean +/- SD) |  | 26.4 +/- 1.7 | 27.4 +/- 1.6 |

### Disease state classification – univariate analysis

Figures 1A-D recapitulates previous analyses by us and others [33, 47–49] showing that Aβ1-42, tTau, pTau-181 and HV are all significantly different in NL and AD subjects and that these values are intermediate for MCI subjects. For all four markers in Figures 1A-D, comparisons of the means between NL and AD groups reveal highly significant differences (p<0.0001 in all cases). However, it should be noted that there is substantial overlap between the distributions in each diagnostic category, rendering these biomarkers unsuitable for use in isolation for diagnostic categorization. As shown in Figures 1E-F, CSF VGF.NSEP levels are depressed in AD patients compared to NL subjects (p=0.0002) and lower levels at baseline are found in future MCI-AD converters than nonconverters (p=0.032).

**Figure 1:**
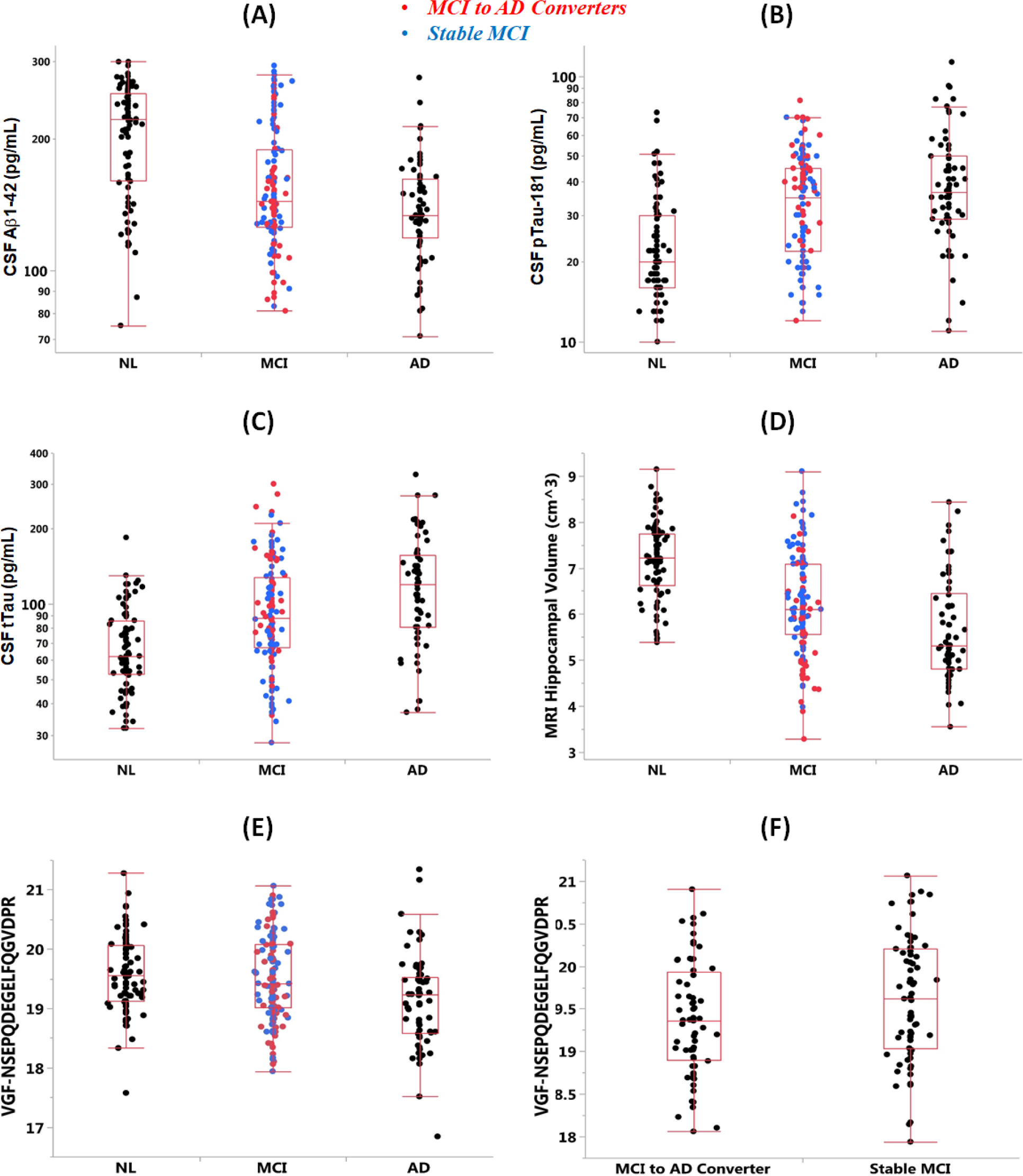
Distributions of biomarkers of in NL, MCI and AD subjects: A) HV, B) Aβ1-42, C) tTau, D) pTau-181, E) VGF.NSEP levels (shown in normalized and log2 transformed intensity units) and F) baseline VGF.NSEP levels in MCI to AD converters and stable MCI subjects over 36 months. In A-E, for the MCI subjects, those that progressed to AD over 36 months are shown in red. The bottom and top ends of the box represent the first and third quartiles respectively, with the line inside the box representing the median. Lines extending out of the ends of the box indicate the range of the data, minus the outliers. The points outside the lines are the low and high outliers.

### Disease state classification – multivariate analysis

To determine if combinations of conventional biomarkers +/− the VGF.NSEP peptide are useful in disease-state classification, data-driven algorithms were used to derive the optimal signature that distinguished NL, MCI and AD. The performances of these signatures are summarized in Table 2. The signatures are grouped into six different categories, as described in the Methods section, and took relatively simple forms. The best performing signature for disease-state classification was a combination of HV + APO-E4 status, with an accuracy of 79.6%. Adding conventional CSF markers (Aβ1-42, tTau and pTau-181 and their ratios) did not enhance this value (accuracy = 76.3%), nor did the addition of VGF.NSEP peptide (accuracy = 75.7%).

**Table 2:** Performance summary of optimal signatures

| Data type | Diagnostic Criteria for Signature positive | AD versus Normal Diagnosis (internal cross-validation) |  |  | 36m MCI Progression to AD (independent validation) |  |  |
| --- | --- | --- | --- | --- | --- | --- | --- |
|  |  | % PPV | % NPV | % Accuracy | % PPV (MCI to AD) | % NPV (Stable MCI) | % Accuracy |
| **AT** | tTau / Ab1-42 > 0.59 | 71.6 | 80.5 | 76.5 | 58.1 | 66.1 | 61.7 |
| **HV** | HV < 6.41 and ApoE4 + | 92.7 | 74.8 | 79.6 | 61.2 | 60.5 | 60.7 |
| **AT + HV** | HV < 7.0, pTau > 18.1, and tTau / Ab1-42 > 0.36 | 73.4 | 78.4 | 76.3 | 60.2 | 70.2 | 64.4 |
| **VGF** | VGF.NSEP < 19.71 and ApoE4 + | **69.1** | **79.1** | **70.4** | **65.9** | **61.5** | **63.0** |
| **AT + VGF** | pTau / Ab1-42 > 0.08, tTau / Ab1-42 > 0.31, and VGF.NSEP < 20.30 | **75.4** | **75.8** | **75.7** | **59.6** | **76.1** | **65.2** |
| **AT + HV + VGF** | HV < 7.81, pTau > 16.18, tTau / Ab1-42 > 0.29, and VGF.NSEP < 20.39 | **72.3** | **78.2** | **75.7** | **62.1** | **79.2** | **68.1** |

### Prediction of the likelihood of MCI to AD progression

As described above, for disease state classification, no advantage was found when adding the VGF.NSEP peptide to the conventional markers (overall accuracy of 76.3% vs. 75.7%, p > 0.05). However, the combined biomarkers signature (HV+AT+VGF) significantly outperformed conventional biomarkers (HV+AT) for the prediction of MCI to AD conversion over 36 months (p=0.00013). Most of the impact of the addition of VGF was in increasing the NPV (from 70.2% to 79.2%, p<0.0001) while the impact on PPV was more modest (60.2% to 62.1%, p=0.008). The signature derived from the conventional and novel markers took a simple form based on only a few markers, with a cut-point on each of them; HV < 7.81 cm^3^, pTau < 16.18 pg/mL, ratio of tTau to Aβ1-42 > 0.29 and VGF.NSEP peptide < 20.39 intensity units. Thus, the addition of a novel VGF peptide to the conventional AD markers provides a simple biomarker that improves the prediction of 36-month disease progression in MCI subjects at baseline.

### Prediction of time to AD progression from MCI

Using available information containing 3-10 year follow-up clinical data, future time to progression was computed using the optimal signatures defined above. Table 3 includes a summary of the median times to progression of the signature negative and signature positive subjects and the overall hazard ratios with 95% confidence intervals. All groups containing conventional biomarkers (combinations of CSF amyloid/tau, HV and APO-E4 status) had similar times to progression (range for 2^nd^ quartile or median = 25.7-31.5 months for signature positive subjects) and hazard ratios (range = 1.9-2.2). By comparison, the signature containing VGF.NSEP + conventional markers performed considerably better with median time to progression of 24.1 months and 96.2 months for the signature positive and signature negative groups respectively, and hazard ratio of 3.9. This difference in hazard ratio is illustrated in Figure 2A (without VGF) and Figure 2B (with VGF), where Kaplan-Meier curves demonstrate time to progression profiles of the signature positive versus signature negative MCI subjects at baseline. The increased separation of the time to progression curves in Figure 2B (with VGF) demonstrates the faster progression experienced by the MCI subjects meeting this signature criterion at baseline.

**Figure 2:**
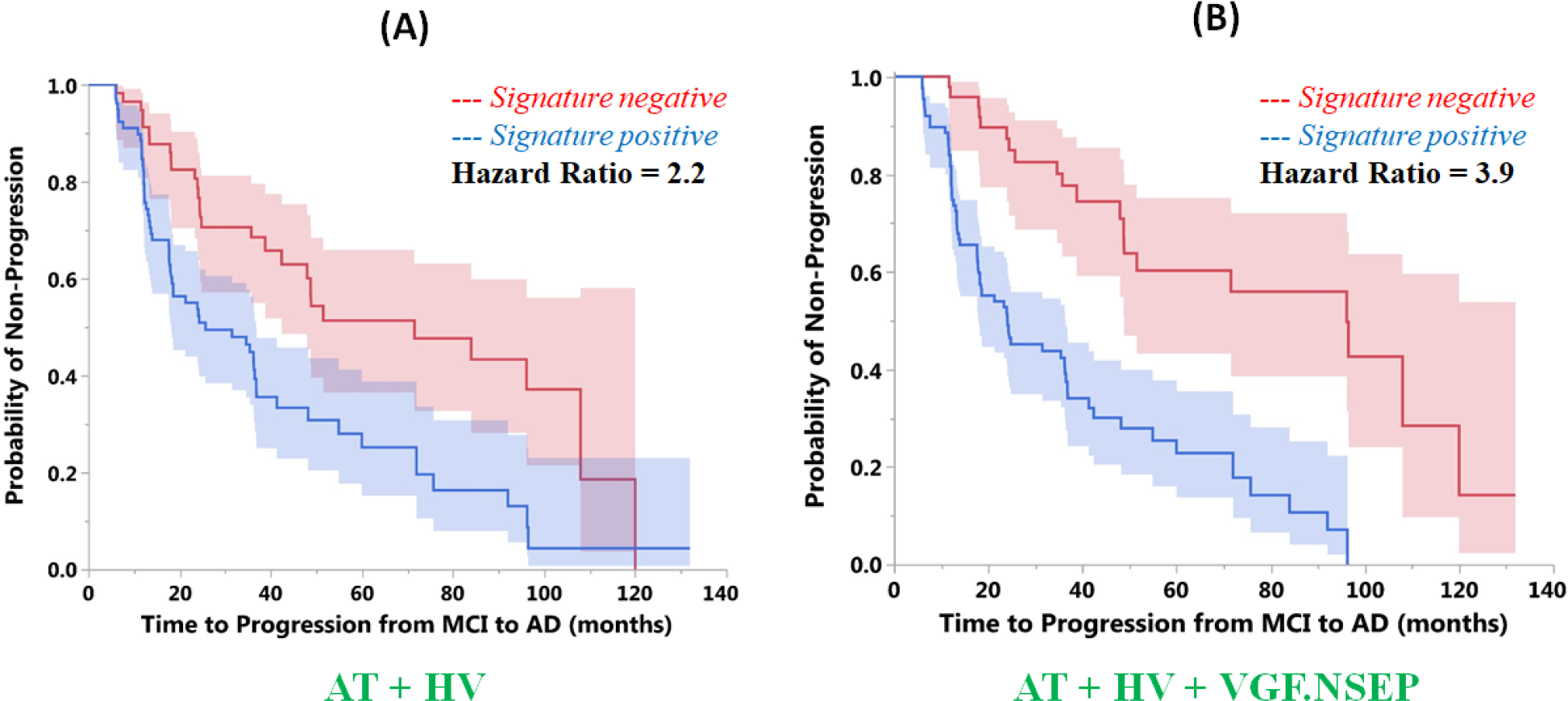
Time to progression profiles of the signature positive versus signature negative MCI subjects with the shaded 95% confidence intervals are shown here via Kaplan-Meier analysis. The effect of signature based on only the conventional markers (HV and AT) is illustrated in Figure 2A and the signature with both the conventional markers and the novel VGF.NSEP peptide from the MRM panel is shown in Figure 2B. Patients meeting the signature criterion that includes the VGF.NSEP peptide experience 3.9-fold faster progression to AD (hazard ratio = 3.9), relative to the 2.2-fold faster progression without this peptide.

**Table 3:**
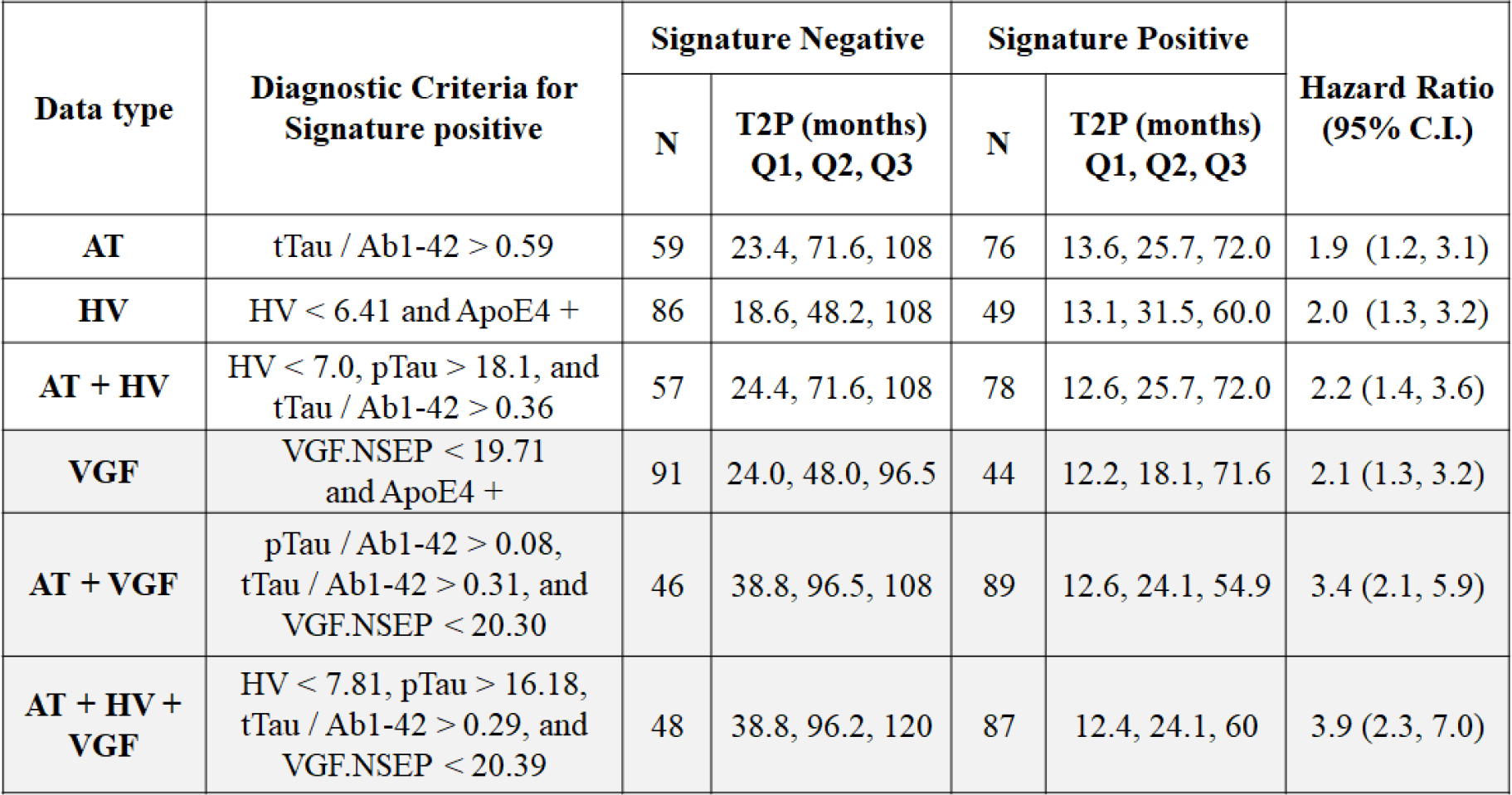
Time to progression (T2P) of MCI subjects to AD using optimal signatures

### Studies of VGF peptide

In further evaluation of the VGF.NSEP peptide, we find that its levels are significantly correlated with pTau-181 and tTau in NL, MCI and AD subjects, and not significantly correlated with Aβ1-42 and brain HV in any of the three groups (see Figures 3A-D). To determine if the impact of VGF was isolated to the particular peptide fragment (VGF.NSEP) that emerged from the multivariate analysis in Llano et al (2017), the other two VGF peptides (AYQGVAAPFPK, referred to as VGF.AYQG and THLGEALAPLSK, referred to as VGF.THLG) in this 320-peptide MRM panel were also assessed. The pairwise correlations are over 97% between the three VGF peptides (Figure 4), and therefore as expected, the other two VGF peptides have very similar effects across the disease states (NL vs. AD significant with p<0.05) and significantly different (p<0.05) between the stable and progressive MCI groups (Figures 5 A-D). When replacing the VGF.NSEP peptide by each of these other two peptides one at a time, the performance of the combined signature for the HV+AT+VGF scenario was quite similar in terms of the median time to progression of MCI subjects to AD (see Table 4 and Figure 6). However, the differences were greater in the overall time course of progression that resulted in larger hazard ratios (4.1 and 4.7). Thus, the considerable improvement we see in the prediction of MCI to AD progression by including VGF with the conventional markers is consistently evident for all three peptide fragments of VGF, and not isolated to a specific peptide fragment.

**Figure 3:**
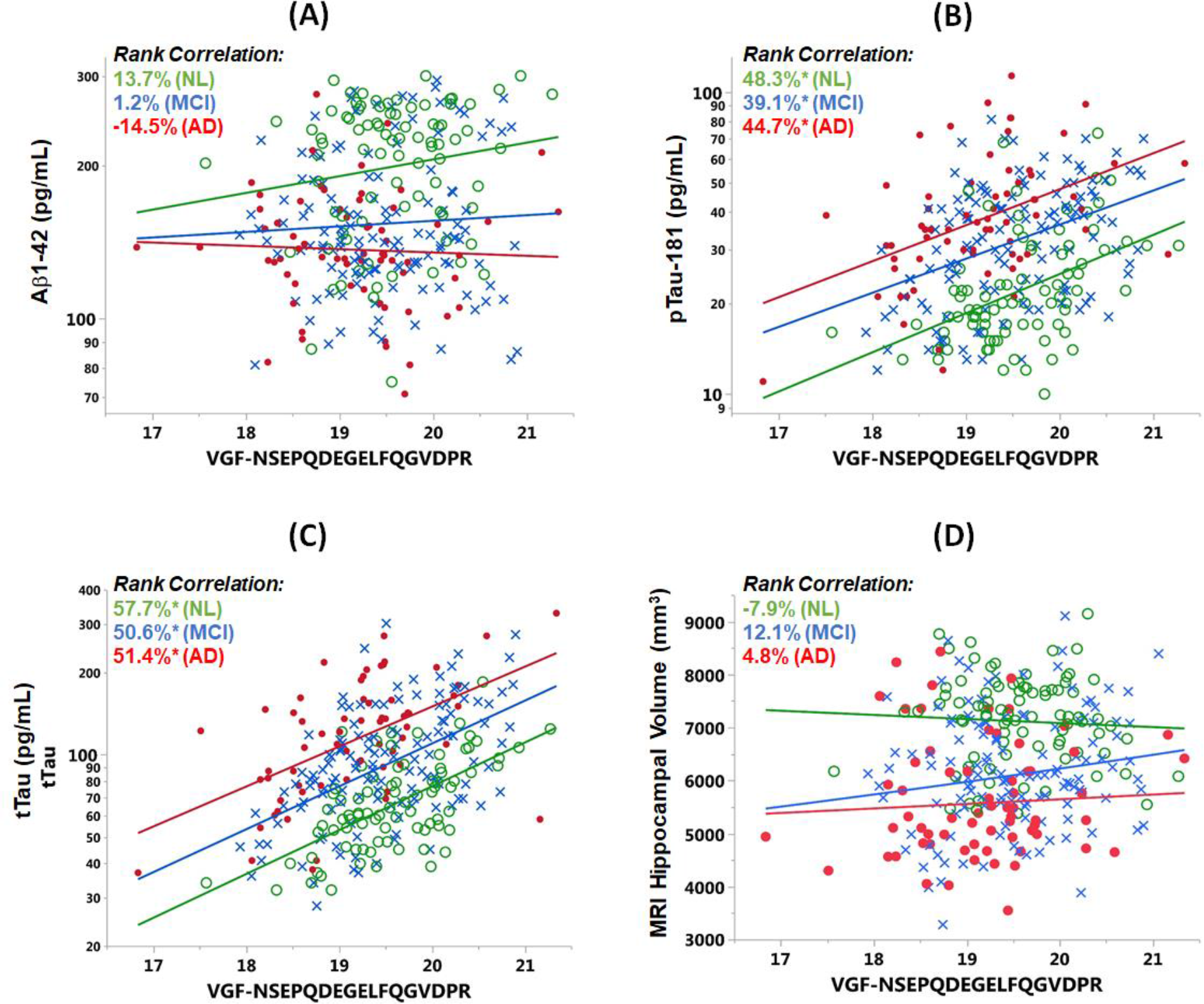
Correlation of the VGF.NSEP peptide levels (shown in normalized and log2 transformed intensity units) versus conventional markers of AD, brain hippocampal volume HV (A), A(1-42 (B), pTau-181 (C), and tTau (D), with the least squares regression lines overlaid on individual subject results from the three groups; Normal (in green), MCI (in blue) and AD (in red). The rank correlation values for each of the groups are shown, with * representing significant correlations (p < 0.05).

**Figure 4:**
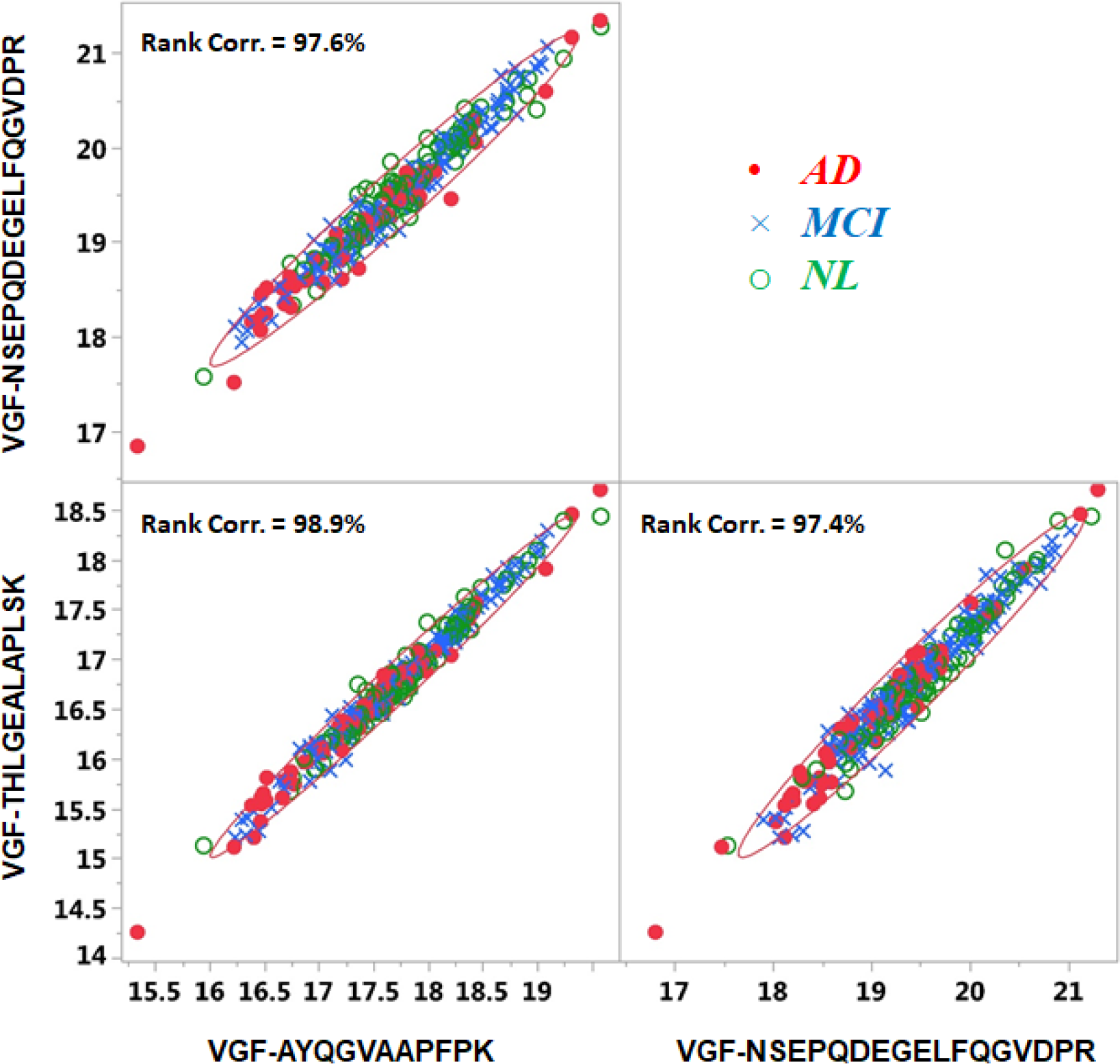
Scatterplot matrix with rank correlation values overlaid for the three VGF peptides levels (shown in normalized and log2 transformed intensity units) from the 320-peptide MRM panel for all subjects.

**Figure 5:**
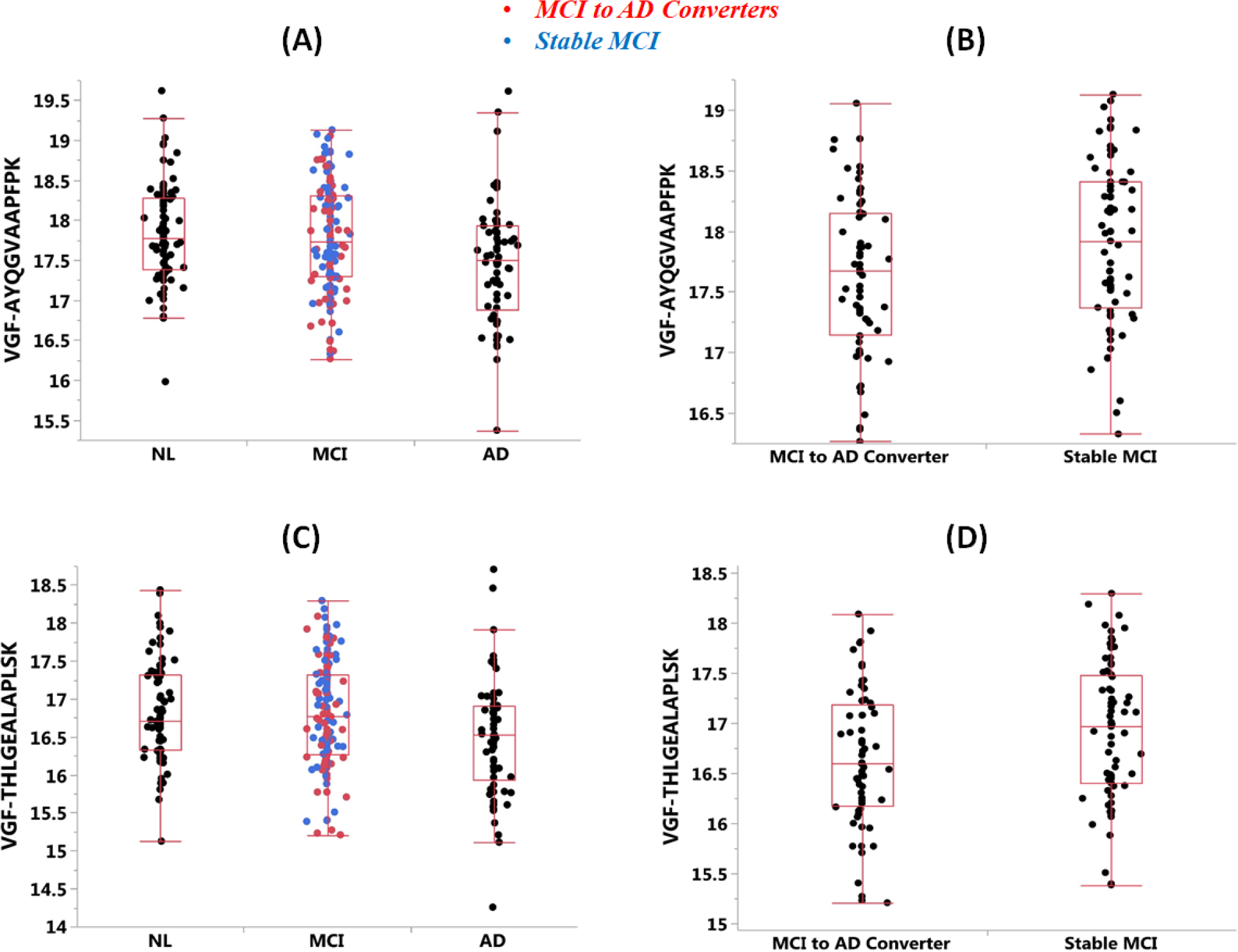
A) Distribution of VGF.AYQG peptide (shown in normalized and log2 transformed intensity units) is shown across the NL, MCI and AD groups, and **B)** among the baseline MCI subjects that either progressed to AD or remained stable over the next 36 months. **C)** Distribution of VGF.THLG peptide (shown in normalized and log2 transformed intensity units) is shown across the NL, MCI and AD groups, and **D)** among the baseline MCI subjects that either progressed to AD or remained stable over the next 36 months.

**Figure 6:**
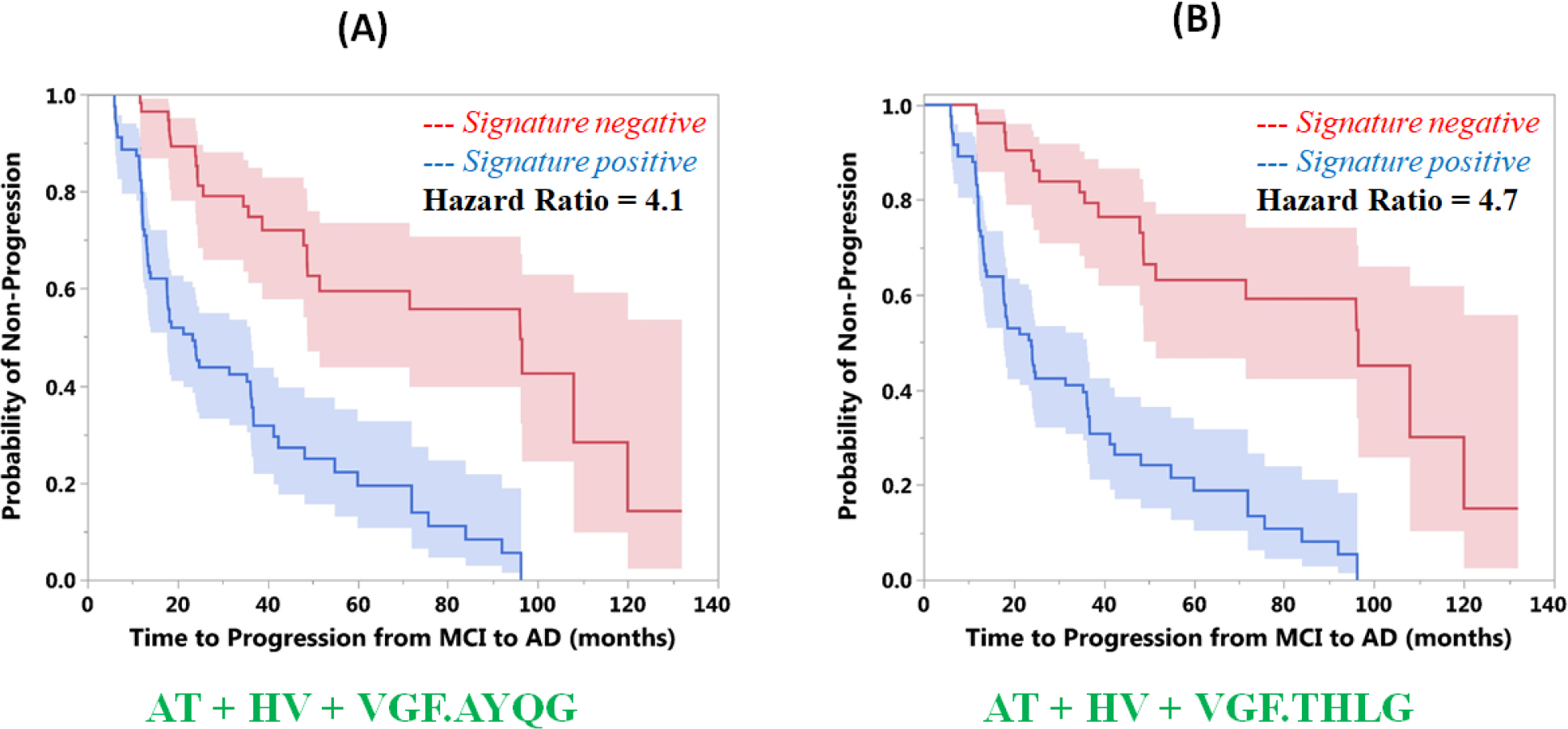
Time to progression profiles for the two additional VGF peptides + conventional biomarkers: (A) AT+HV+VGF.AYQG and (B) AT+HV+VGF.THLG. In both cases, the signature positive versus signature negative MCI subjects with the shaded 95% confidence intervals are shown via Kaplan-Meier analysis.

**Table 4:** Time to progression (T2P) of MCI subjects to AD using optimal and other candidate signatures for the AT+HV+MRM scenario

| Data type | Diagnostic Criteria for Signature positive | Signature Negative |  | Signature Positive |  | Hazard Ratio (95% C.I.) |
| --- | --- | --- | --- | --- | --- | --- |
|  |  | N | T2P (months)<br>Q1, Q2, Q3 | N | T2P (months)<br>Q1, Q2, Q3 |  |
| AT + HV + VGF | HV < 7.81, pTau > 16.18, tTau / Ab1-42 > 0.29, and VGF.NSEP < 20.39 | 48 | 38.8, 96.2, 120 | 87 | 12.4, 24.1, 60.0 | 3.9 (2.3, 7.0) |
|  | HV < 7.81, pTau > 16.18, tTau / Ab1-42 > 0.29, and VGF.AYQG < 18.47 | 56 | 35.8, 96.2, 120 | 79 | 12.3, 23.4, 48.3 | 4.1 (2.4, 6.8) |
|  | HV < 7.81, pTau > 16.18, tTau / Ab1-42 > 0.29, and VGF.THLG < 17.62 | 52 | 48.0, 96.5, 120 | 83 | 12.3, 23.9, 48.3 | 4.7 (2.7, 8.2) |

## Discussion

### Summary

Here, we examined the ability of CSF VGF-derived peptides, in combination with conventional AD biomarkers (Aβ1-42, tTau, pTau-181, their ratios and HV) to serve as a disease-state marker to distinguish between AD and NLn subjects, and to predict conversion from MCI to AD in a separate group of subjects. We observed that CSF levels of a VGF peptide, on its own, are lower in AD subjects than NLs and that lower levels predict MCI to AD conversion. When combined with conventional biomarkers, the VGF peptide significantly increased the ability of a combination of conventional biomarkers to predict MCI to AD conversion, with the hazard ratio increasing from 2.2 to 3.9. These data suggest that VGF may play a previously under-recognized role in the pathophysiology of AD and that CSF VGF may be useful to help predict MCI to AD conversion.

### Total tau vs. phosphorylated tau in predicting MCI to AD conversion

It is notable that, when combined with HV, Aβ1-42 and VGF.NSEP, CSF tTau was found to more strongly predict MCI to AD conversion than pTau-181. tTau, but not pTau-181, elevations in the CSF have been observed in many non-AD conditions involving neuronal injury, including stroke, traumatic brain injury, Creutzfeldt-Jacob disease, multiple sclerosis as well as vascular dementia [50–55], suggesting that tTau is a general marker of neuronal injury, while pTau-181 better reflects AD pathology. The finding in the current study that tTau is more strongly predictive of MCI-AD conversion than pTau-181 is consistent with previous data showing that total tau is more predictive than pTau-181 in predicting subsequent cognitive decline in MCI and AD [56, 57]. These findings suggest that while pTau-181 may be more useful as a disease-state marker, particularly when making a differential diagnosis, that tTau may be a better marker of disease activity and thus the current rate of clinical decline. In addition, because the database we used only captures the progression to AD of these MCI subjects, and not the other neurodegenerative diseases, is likely that the use of pTau-181 instead of tTau in our signature may have shown improved performance specificity if we had applied it to a broader group of MCI subjects that also experienced progression to the other forms of dementia.

### VGF and AD

The current finding that all peptides associated with VGF are diminished in the CSF of AD patients compared to controls is consistent with multiple previous studies comparing VGF peptide or protein levels in CSF [26–30, 32] and brain tissue (parietal cortex [22]) from AD and control subjects. The functional significance of this decrease is not yet clear but may relate to VGF’s potential role in synaptic plasticity and/or neuronal metabolism. VGF is found widely throughout the brain, including areas highly affected in AD such as cerebral cortex, hippocampus, entorhinal cortex, basal forebrain, amygdala, and brainstem [22, 58, 59]. Its expression is upregulated by neuronal activity [60] and can be induced by neuronal growth factors such as brain-derived neurotrophic factor (BDNF [58, 61]). In animal models, VGF has been shown to be important for the mediation of synaptic plasticity and neurogenesis in the hippocampus [58, 61–63], and knock out of this gene has been shown to cause significant anorexia [64], while overexpression may protect the brain against AD-related pathology [21]. These functions align well with the loss of hippocampal function and significant anorexia seen in AD [65, 66].

The mechanism behind the drop in VGF levels in AD CSF is not yet clear. Given the parallel drop in the cerebral cortex [22], low levels in the CSF are likely not due to a shift of VGF from CSF to parenchyma, as has been hypothesized for the low levels of Aβ in the CSF of AD patients [67]. Low levels of VGF in CSF (and brain) may suggest that VGF is a general marker for neuronal loss, consistent the drop in CSF VGF in frontotemporal dementia [68], as such, potentially putting VGF into the “neurodegenerative/neuronal injury” class of biomarkers in the AT(N) framework previously described [69]. This notion that low CSF VGF may be a reflection of neuronal damage is consistent with the current data which demonstrate that VGF levels are correlated with hippocampal volume as well as tTau and pTau-181 levels (Figure 3). Future work examining VGF across other states of neuronal injury may help to add clarity to this issue.

One previous study observed borderline elevations of VGF in the CSF of MCI compared to control and AD subjects, and that VGF elevations in MCI subjects predicted later conversion to AD [32]. Such transient elevations are reminiscent of “pseudonormalization” of other biomarkers whose values in MCI appear to change in the opposite direction of that seen in AD [20, 34, 70]. It is not clear from the Duits et al. report which specific peptides were elevated in MCI, though the two peptides examined in their study (NSEPQDEGELFQGVDPR and AYQGVAAPFPK) matched two of the three peptides in the current study, all of which showed decreases in MCI and AD (Figure 5). The source of the apparent discrepancy is not yet clear, though we note that all analytes, not just VGF, in the Duits et al. study showed elevations in the MCI group. It is notable that other analytes that are elevated in MCI subjects in the Duits et al. study such as Chromogranin A have been found to be unchanged in other studies [71] or, in the case of VGF, decreased in MCI patients that convert to AD [27], suggesting a more general difference in the databases or the analytical methodologies used between the Duits et al. study and other studies.

### Implications of the prediction of MCI-AD conversion

CSF Aβ1-42 and tau derivatives as biomarkers are well-established for the prediction of clinical decline in MCI [72–76] (for meta-analyses see [77, 78]). In addition, predictive accuracy of these markers increases when they are combined with volumetric imaging markers [79–83]. Both of these findings were reproduced in the current study (Table 2). In addition, recently a number of non-Aβ, non-tau CSF markers have been found, often using proteomic approaches, that separate AD from NL subjects, and these markers have been implicated across a number of metabolic, inflammatory and synaptic physiology pathways [25–29, 31, 84–90]. A small number have also shown the ability to predict MCI to AD conversion. For example, heart fatty acid binding protein, chemokine receptor 2, neurogranin, calbindin, IL-1, thymus-expressed chemokine have all individually been shown to predict MCI to AD progression [14–20]. In addition, we and others identified panels of peptides that predict MCI to AD progression [19, 20]. These data point to a range of potential pathophysiological mechanisms implicated in AD outside of the classical amyloid-driven cascade. It will be important to replicate the findings in this study as well as others in independent cohorts. In addition, like most of the previous work, the current study did not examine non-AD dementia or other neurologic disease. This absence is particularly important in the current study which shows VGF levels that correlate with tTau levels (a marker of neurodegeneration, as described above) but not hippocampal volume (Figures 3C and D). These data suggest that VGF levels may correlate with a more general neurodegenerative phenotype. Therefore, it will be important in future studies to include non-AD dementias as well as other neurological illness such as stroke or encephalitis, to determine the specificity of VGF as a biomarker for AD and predictor of MCI to AD progression.

## Acknowledgments

The authors thank Professor Danielle Harvey from UC Davis for providing valuable input regarding the ADNI imaging data.

## Notes

Disclosures: Viswanath Devanarayan is an employee of Charles River Laboratories, and as such owns equity in, receives salary and other compensation from Charles River Laboratories.

## References

1. Dubois, B., et al., Preclinical Alzheimer’s disease: definition, natural history, and diagnostic criteria. Alzheimer’s & Dementia, 2016. 12(3): p. 292–323.

2. Bloom, G.S., Amyloid-β and tau: the trigger and bullet in Alzheimer disease pathogenesis. JAMA neurology, 2014. 71(4): p. 505–508.

3. Haass, C. and D.J. Selkoe, Soluble protein oligomers in neurodegeneration: lessons from the Alzheimer’s amyloid β-peptide. Nature reviews Molecular cell biology, 2007. 8(2): p. 101.

4. Sengupta, U., A.N. Nilson, and R. Kayed, The role of amyloid-β oligomers in toxicity, propagation, and immunotherapy. EBioMedicine, 2016. 6: p. 42–49.

5. Blennow, K., et al., Cerebrospinal fluid and plasma biomarkers in Alzheimer disease. Nature Reviews Neurology, 2010. 6(3): p. 131.

6. Perrin, R.J., A.M. Fagan, and D.M. Holtzman, Multimodal techniques for diagnosis and prognosis of Alzheimer’s disease. Nature, 2009. 461(7266): p. 916.

7. Dubois, B., et al., Research criteria for the diagnosis of Alzheimer’s disease: revising the NINCDS–ADRDA criteria. The Lancet Neurology, 2007. 6(8): p. 734–746.

8. Rasmusson, D., et al., Accuracy of clinical diagnosis of Alzheimer disease and clinical features of patients with non-Alzheimer disease neuropathology. Alzheimer disease and associated disorders, 1996. 10(4): p. 180–188.

9. Kirson, N., et al., Excess costs associated with misdiagnosis of Alzheimer’s disease among US Medicare beneficiaries with vascular dementia or Parkinson’s disease. Alzheimer’s & Dementia: The Journal of the Alzheimer’s Association, 2013. 9(4): p. P845–P846.

10. Beach, T.G., et al., Accuracy of the clinical diagnosis of Alzheimer disease at National Institute on Aging Alzheimer Disease Centers, 2005–2010. Journal of neuropathology and experimental neurology, 2012. 71(4): p. 266–273.

11. Qian, W., et al., Misdiagnosis of Alzheimer’s disease: Inconsistencies between clinical diagnosis and neuropathological confirmation. Alzheimer’s & Dementia: The Journal of the Alzheimer’s Association, 2016. 12(7): p. P293.

12. Yu, P., et al., Enriching amnestic mild cognitive impairment populations for clinical trials: optimal combination of biomarkers to predict conversion to dementia. Journal of Alzheimer’s disease, 2012. 32(2): p. 373–385.

13. van Rossum, I.A., et al., Biomarkers as predictors for conversion from mild cognitive impairment to Alzheimer-type dementia: implications for trial design. Journal of Alzheimer’s Disease, 2010. 20(3): p. 881–891.

14. Westin, K., et al., CCL2 is associated with a faster rate of cognitive decline during early stages of Alzheimer’s disease. PloS one, 2012. 7(1): p. e30525.

15. Chiasserini, D., et al., CSF levels of heart fatty acid binding protein are altered during early phases of Alzheimer’s disease. Journal of Alzheimer’s Disease, 2010. 22(4): p. 1281–1288.

16. Olsson, B., et al., Cerebrospinal fluid levels of heart fatty acid binding protein are elevated prodromally in Alzheimer’s disease and vascular dementia. Journal of Alzheimer’s Disease, 2013. 34(3): p. 673–679.

17. Kester, M.I., et al., Neurogranin as a cerebrospinal fluid biomarker for synaptic loss in symptomatic Alzheimer disease. JAMA neurology, 2015. 72(11): p. 1275–1280.

18. Craig-Schapiro, R., et al., Multiplexed immunoassay panel identifies novel CSF biomarkers for Alzheimer’s disease diagnosis and prognosis. PloS one, 2011. 6(4): p. e18850.

19. Spellman, D.S., et al., Development and evaluation of a multiplexed mass spectrometry based assay for measuring candidate peptide biomarkers in Alzheimer’s Disease Neuroimaging Initiative (ADNI) CSF. Proteomics-Clinical Applications, 2015.

20. Llano, D.A., et al., A multivariate predictive modeling approach reveals a novel CSF peptide signature for both Alzheimer’s Disease state classification and for predicting future disease progression. PloS one, 2017. 12(8): p. e0182098.

21. Beckmann, N.D., et al., Multiscale causal network models of Alzheimer’s disease identify VGF as a key regulator of disease. bioRxiv, 2018.

22. Cocco, C., et al., Distribution of VGF peptides in the human cortex and their selective changes in Parkinson’s and Alzheimer’s diseases. Journal of anatomy, 2010. 217(6): p. 683–693.

23. Levi, A., et al., Processing, distribution, and function of VGF, a neuronal and endocrine peptide precursor. Cellular and molecular neurobiology, 2004. 24(4): p. 517–533.

24. Salton, S.R., et al., VGF: a novel role for this neuronal and neuroendocrine polypeptide in the regulation of energy balance. Frontiers in neuroendocrinology, 2000. 21(3): p. 199–219.

25. Wijte, D., et al., A novel peptidomics approach to detect markers of Alzheimer’s disease in cerebrospinal fluid. Methods, 2012. 56(4): p. 500–507.

26. Hendrickson, R.C., et al., High resolution discovery proteomics reveals candidate disease progression markers of Alzheimer’s disease in human cerebrospinal fluid. PloS one, 2015. 10(8): p. e0135365.

27. Jahn, H., et al., Peptide fingerprinting of Alzheimer’s disease in cerebrospinal fluid: identification and prospective evaluation of new synaptic biomarkers. PloS one, 2011. 6(10): p. e26540.

28. Selle, H., et al., Identification of novel biomarker candidates by differential peptidomics analysis of cerebrospinal fluid in Alzheimer’s disease. Combinatorial chemistry & high throughput screening, 2005. 8(8): p. 801–806.

29. Hoölttä, M., et al., An Integrated Workflow for Multiplex CSF Proteomics and Peptidomics Identification of Candidate Cerebrospinal Fluid Biomarkers of Alzheimer’s Disease. Journal of proteome research, 2014. 14(2): p. 654–663.

30. Carrette, O., et al., A panel of cerebrospinal fluid potential biomarkers for the diagnosis of Alzheimer’s disease. PROTEOMICS: International Edition, 2003. 3(8): p. 1486–1494.

31. Brinkmalm, G., et al., A parallel reaction monitoring mass spectrometric method for analysis of potential CSF biomarkers for Alzheimer’s disease. PROTEOMICS–Clinical Applications, 2018. 12(1): p. 1700131.

32. Duits, F.H., et al., Synaptic proteins in CSF as potential novel biomarkers for prognosis in prodromal Alzheimer’s disease. Alzheimer’s research & therapy, 2018. 10(1): p. 5.

33. Devanarayan, P., et al., Identification of a simple and novel cut-point based CSF and MRI signature for predicting Alzheimer’s disease progression that reinforces the 2018 NIA-AA research framework Journal of Alzheimer’s Disease, 2019. In Press.

34. Llano, D.A., et al., Evaluation of plasma proteomic data for Alzheimer disease state classification and for the prediction of progression from mild cognitive impairment to Alzheimer disease. Alzheimer Disease & Associated Disorders, 2013. 27(3): p. 233–243.

35. Jack, C.R., et al., Prediction of AD with MRI-based hippocampal volume in mild cognitive impairment. Neurology, 1999. 52(7): p. 1397–1397.

36. Risacher, S.L., et al., Baseline MRI predictors of conversion from MCI to probable AD in the ADNI cohort. Current Alzheimer Research, 2009. 6(4): p. 347–361.

37. Dubois, B., et al., Revising the definition of Alzheimer’s disease: a new lexicon. The Lancet Neurology, 2010. 9(11): p. 1118–1127.

38. Dale, A.M., B. Fischl, and M.I. Sereno, Cortical surface-based analysis: I. Segmentation and surface reconstruction. Neuroimage, 1999. 9(2): p. 179–194.

39. Fischl, B., M.I. Sereno, and A.M. Dale, Cortical surface-based analysis: II: inflation, flattening, and a surface-based coordinate system. Neuroimage, 1999. 9(2): p. 195–207.

40. Fischl, B., et al., High-resolution intersubject averaging and a coordinate system for the cortical surface. Human brain mapping, 1999. 8(4): p. 272–284.

41. Shaw, L.M., et al., Cerebrospinal fluid biomarker signature in Alzheimer’s disease neuroimaging initiative subjects. Annals of neurology, 2009. 65(4): p. 403–413.

42. Chen, G., et al., A PRIM approach to predictive-signature development for patient stratification. Statistics in medicine, 2015. 34(2): p. 317–342.

43. Huang, X., et al., Patient subgroup identification for clinical drug development. Statistics in medicine, 2017. 36(9): p. 1414–1428.

44. Corder, E., et al., Gene dose of apolipoprotein E type 4 allele and the risk of Alzheimer’s disease in late onset families. Science, 1993. 261(5123): p. 921–923.

45. Drzezga, A., et al., Prediction of individual clinical outcome in MCI by means of genetic assessment and 18F-FDG PET. Journal of Nuclear Medicine, 2005. 46(10): p. 1625–1632.

46. Lindsay, J., et al., Risk factors for Alzheimer’s disease: a prospective analysis from the Canadian Study of Health and Aging. American journal of epidemiology, 2002. 156(5): p. 445–453.

47. Galasko, D., et al., High cerebrospinal fluid tau and low amyloid β42 levels in the clinical diagnosis of Alzheimer disease and relation to apolipoprotein E genotype. Archives of neurology, 1998. 55(7): p. 937–945.

48. Tapiola, T., et al., Cerebrospinal fluid β-amyloid 42 and tau proteins as biomarkers of Alzheimer-type pathologic changes in the brain. Archives of neurology, 2009. 66(3): p. 382–389.

49. Hampel, H., et al., Value of CSF β-amyloid 1–42 and tau as predictors of Alzheimer’s disease in patients with mild cognitive impairment. Molecular psychiatry, 2004. 9(7): p. 705.

50. Satoh, K., et al., 14-3-3 protein, total tau and phosphorylated tau in cerebrospinal fluid of patients with Creutzfeldt-Jakob disease and neurodegenerative disease in Japan. Cellular and molecular neurobiology, 2006. 26(1): p. 45–52.

51. Nägga, K., et al., Cerebrospinal fluid phospho-Tau, total Tau and β-Amyloid1–42 in the differentiation between Alzheimer’s disease and vascular dementia. Dementia and geriatric cognitive disorders, 2002. 14(4): p. 183–190.

52. Kapaki, E., et al., Cerebrospinal fluid tau, phospho-tau181 and β-amyloid1-42 in idiopathic normal pressure hydrocephalus: a discrimination from Alzheimer’s disease. European journal of neurology, 2007. 14(2): p. 168–173.

53. Skillbäck, T., et al., Diagnostic performance of cerebrospinal fluid total tau and phosphorylated tau in Creutzfeldt-Jakob disease: results from the Swedish Mortality Registry. JAMA neurology, 2014. 71(4): p. 476–483.

54. Hesse, C., et al., Transient increase in total tau but not phospho-tau in human cerebrospinal fluid after acute stroke. Neuroscience letters, 2001. 297(3): p. 187–190.

55. Bartosik-Psujek, H. and Z. Stelmasiak, The CSF levels of total-tau and phosphotau in patients with relapsing-remitting multiple sclerosis. Journal of neural transmission, 2006. 113(3): p. 339–345.

56. SämgÃ¥rd, K., et al., Cerebrospinal fluid total tau as a marker of Alzheimer’s disease intensity. International Journal of Geriatric Psychiatry: A journal of the psychiatry of late life and allied sciences, 2010. 25(4): p. 403–410.

57. Blom, E.S., et al., Rapid progression from mild cognitive impairment to Alzheimer’s disease in subjects with elevated levels of tau in cerebrospinal fluid and the APOE ε4/ε4 genotype. Dementia and geriatric cognitive disorders, 2009. 27(5): p. 458–464.

58. Alder, J., et al., Brain-derived neurotrophic factor-induced gene expression reveals novel actions of VGF in hippocampal synaptic plasticity. Journal of Neuroscience, 2003. 23(34): p. 10800–10808.

59. Snyder, S.E. and S.R. Salton, Expression of VGF mRNA in the adult rat central nervous system. Journal of Comparative Neurology, 1998. 394(1): p. 91–105.

60. Snyder, S., et al., The messenger RNA encoding VGF, a neuronal peptide precursor, is rapidly regulated in the rat central nervous system by neuronal activity, seizure and lesion. Neuroscience, 1997. 82(1): p. 7–19.

61. Thakker-Varia, S., et al., The neuropeptide VGF produces antidepressant-like behavioral effects and enhances proliferation in the hippocampus. Journal of Neuroscience, 2007. 27(45): p. 12156–12167.

62. Bozdagi, O., et al., The neurotrophin-inducible gene Vgf regulates hippocampal function and behavior through a brain-derived neurotrophic factor-dependent mechanism. Journal of Neuroscience, 2008. 28(39): p. 9857–9869.

63. Thakker-Varia, S., et al., VGF (TLQP-62)-induced neurogenesis targets early phase neural progenitor cells in the adult hippocampus and requires glutamate and BDNF signaling. Stem cell research, 2014. 12(3): p. 762–777.

64. Hahm, S., et al., Targeted deletion of the Vgf gene indicates that the encoded secretory peptide precursor plays a novel role in the regulation of energy balance. Neuron, 1999. 23(3): p. 537–548.

65. Gillette-Guyonnet, S., et al., Weight loss in Alzheimer disease. The American journal of clinical nutrition, 2000. 71(2): p. 637S–642S.

66. Johnson, D.K., C.H. Wilkins, and J.C. Morris, Accelerated weight loss may precede diagnosis in Alzheimer disease. Archives of neurology, 2006. 63(9): p. 1312–1317.

67. Motter, n., et al., Reduction of β-amyloid peptide42 in the cerebrospinal fluid of patients with Alzheimer’s disease. Annals of Neurology: Official Journal of the American Neurological Association and the Child Neurology Society, 1995. 38(4): p. 643–648.

68. Rüetschi, U., et al., Identification of CSF biomarkers for frontotemporal dementia using SELDI-TOF. Experimental neurology, 2005. 196(2): p. 273–281.

69. Jack, C.R., et al., NIA-AA Research Framework: Toward a biological definition of Alzheimer’s disease. Alzheimer’s & Dementia, 2018. 14(4): p. 535–562.

70. Dickerson, B., et al., Increased hippocampal activation in mild cognitive impairment compared to normal aging and AD. Neurology, 2005. 65(3): p. 404–11.

71. Perrin, R.J., et al., Identification and validation of novel cerebrospinal fluid biomarkers for staging early Alzheimer’s disease. PloS one, 2011. 6(1): p. e16032.

72. Hansson, O., et al., Association between CSF biomarkers and incipient Alzheimer’s disease in patients with mild cognitive impairment: a follow-up study. The Lancet Neurology, 2006. 5(3): p. 228–234.

73. Forlenza, O.V., et al., Cerebrospinal fluid biomarkers in Alzheimer’s disease: Diagnostic accuracy and prediction of dementia. Alzheimer’s & Dementia: Diagnosis, Assessment & Disease Monitoring, 2015. 1(4): p. 455–463.

74. Zetterberg, H., L.-O. Wahlund, and K. Blennow, Cerebrospinal fluid markers for prediction of Alzheimer’s disease. Neuroscience letters, 2003. 352(1): p. 67–69.

75. Handels, R.L., et al., Predicting progression to dementia in persons with mild cognitive impairment using cerebrospinal fluid markers. Alzheimer’s & Dementia, 2017. 13(8): p. 903–912.

76. Lanari, A. and L. Parnetti, Cerebrospinal fluid biomarkers and prediction of conversion in patients with mild cognitive impairment: 4-year follow-up in a routine clinical setting. The Scientific World Journal, 2009. 9: p. 961–966.

77. Diniz, B.S., J.A. Pinto Jr, and O.V. Forlenza, Do CSF total tau, phosphorylated tau, and β-amyloid 42 help to predict progression of mild cognitive impairment to Alzheimer’s disease? A systematic review and meta-analysis of the literature. The World Journal of Biological Psychiatry, 2008. 9(3): p. 172–182.

78. Mitchell, A.J., CSF phosphorylated tau in the diagnosis and prognosis of mild cognitive impairment and Alzheimer’s disease–a meta-analysis of 51 studies. Journal of Neurology, Neurosurgery & Psychiatry, 2009.

79. Vemuri, P., et al., MRI and CSF biomarkers in normal, MCI, and AD subjects: predicting future clinical change. Neurology, 2009. 73(4): p. 294–301.

80. Nesteruk, M., et al., Combined use of biochemical and volumetric biomarkers to assess the risk of conversion of mild cognitive impairment to Alzheimer’s disease. Folia neuropathologica, 2016. 54(4): p. 369–374.

81. Bouwman, F., et al., CSF biomarkers and medial temporal lobe atrophy predict dementia in mild cognitive impairment. Neurobiology of aging, 2007. 28(7): p. 1070–1074.

82. Heister, D., et al., Predicting MCI outcome with clinically available MRI and CSF biomarkers. Neurology, 2011: p. WNL. 0b013e3182343314.

83. Westman, E., J.-S. Muehlboeck, and A. Simmons, Combining MRI and CSF measures for classification of Alzheimer’s disease and prediction of mild cognitive impairment conversion. Neuroimage, 2012. 62(1): p. 229–238.

84. Maarouf, C.L., et al., Proteomic analysis of Alzheimer’s disease cerebrospinal fluid from neuropathologically diagnosed subjects. Current Alzheimer research, 2009. 6(4): p. 399–406.

85. Oláh, Z., et al., Proteomic analysis of cerebrospinal fluid in Alzheimer’s disease: wanted dead or alive. Journal of Alzheimer’s Disease, 2015. 44(4): p. 1303–1312.

86. Roher, A.E., et al., Proteomics-derived cerebrospinal fluid markers of autopsy-confirmed Alzheimer’s disease. Biomarkers, 2009. 14(7): p. 493–501.

87. Choi, Y.S., et al., Targeted human cerebrospinal fluid proteomics for the validation of multiple Alzheimer’s disease biomarker candidates. Journal of Chromatography B, 2013. 930: p. 129–135.

88. Wildsmith, K.R., et al., Identification of longitudinally dynamic biomarkers in Alzheimer’s disease cerebrospinal fluid by targeted proteomics. Molecular neurodegeneration, 2014. 9(1): p. 22.

89. Paterson, R., et al., A targeted proteomic multiplex CSF assay identifies increased malate dehydrogenase and other neurodegenerative biomarkers in individuals with Alzheimer’s disease pathology. Translational psychiatry, 2016. 6(11): p. e952.

90. Heywood, W.E., et al., Identification of novel CSF biomarkers for neurodegeneration and their validation by a high-throughput multiplexed targeted proteomic assay. Molecular neurodegeneration, 2015. 10(1): p. 64.

